# skiftiTools: An R package for reading, writing, analysing, and visualising, tract-based spatial statistics (TBSS) derived diffusion MR images

**DOI:** 10.1101/2025.10.26.684698

**Authors:** Jetro J. Tuulari, Aaron Barron, Ashmeet Jolly, Ilkka Suuronen, Hilyatushalihah K. Audah, Aylin Rosberg, Isabella L.C. Mariani Wigley, Elena Vartiainen, Silja Luotonen, Elmo P. Pulli, Hasse Karlsson, Riikka Korja, Linnea Karlsson, Antti Airola, Jakob Seidlitz, Richard A.I. Bethlehem, Harri A. Merisaari

**Author notes:** **Corresponding author:** Jetro J. Tuulari,. FinnBrain study, Medisiina A3, Kiinanmyllynkatu 4-8, 20540, Turku, Finland.

## Abstract

skiftiTools processes three- and four-dimensional neuroimaging data, facilitating advanced statistical modelling with voxelwise data in any software of choice. Tract-Based Spatial Statistics (TBSS) is a conventionally used tool to make statistical calculations in voxel space for brain imaging data. While pre-existing software packages provide support for general linear model based statistics, there is a clear need for more sophisticated modeling. skiftiTools writes subject-per-volume NIfTI files as tab-separated value ASCII files, which are easily readable by most commonly used statistical tools such as R language (RStudio), SPSS, SAS, and GraphPad Prism. This facilitates a wide range of voxel-level statistical analyses from TBSS data, including estimation of standardised effect sizes, clustering, dimensionality reduction, non-linear and machine learning predictive modelling, which we showcase in this article using FinnBrain and developing Human Connectome Project diffusion MRI data. After statistical processing, the resulting ASCII data can then be read again for visualization. The package supports NIfTI image format, tab-separated ASCII format, and its own stand-alone format for efficient disk usage. It is open source (https://github.com/haanme/skiftiTools), built on R-language and has easy installation from R’s CRAN package repository. In addition, we provide basic functions available in Docker containers for further platform independence.

**Highlights:**

- The skiftiTools R package is an open-source, user-friendly interface for analysing voxelwise diffusion tensor imaging (DTI) data following tract-based spatial statistics (TBSS) processing
- It supports reading, writing, visualization, mathematical operations, and data manipulation and thus allows comprehensive conventional and advanced statistics, including machine learning
- skiftiTools bridges a critical gap between statistical tools in R and voxelwise neuroimaging data – including comparable means to perform multiple comparison corrections and much needed possibility to use non-linear statistics

## 1. Introduction

With increasing availability of large scale neuroimaging data, there is dire need for integrated statistical applications. It has become essential to produce end-to-end pipelines/workflows that can bridge the gap between raw image formatting of neuroimaging data to common and evolving statistical software tools. In this context, it is not surprising that there has been increased interest and effort in bridging the tabular image file formats to statistical applications ^1–7^. One of the most pertinent general limitations to contemporary neuroscience is that statistics are focused on linear modeling, and available tools do not offer easy to use means to model non-linear associations, even though they are known to be common. Existing tools similarly have significant limitations with flexible use of statistics including reporting of effect sizes as well as use of machine learning and often lack spatially informed multiple comparison corrections that are routinely done in image-based statistics. We address these limitations in the current article with the creation and implementation of a novel R software-based solution called skiftiTools.

Prior work bridging neuroimaging and R software ^1–7^ has mainly focused on manipulations of CIFTI files pertaining to structural magnetic resonance imaging (MRI) measures mapped across the cortical surface ^8^. These approaches are welcome and justified since they have the potential to provide easy access to work with neuroimaging data from a variety of disciplines with markedly less technical expertise in neuroimaging tools, and can provide easier approaches to education ^4,6^. Existing tools allow mapping the MRI data between image and tabular formats for the cerebral cortex, subcortical structures, and the cerebellum. Importantly, diffusion tensor imaging (DTI) ^9^, a popular technique to study the human white matter, remains largely without similar standard data formats and analysis applications. When researchers wish to apply statistics to neuroimaging data, they are generally confronted with two options: either well-established software that offer general linear models but limited flexibility, or the requirement to develop new ‘in-house’ tools, often requiring high-level expertise in programming and statistics. This is especially the case for tract-based spatial statistics (TBSS) data ^10^, where current tools do not offer compatibility with advanced statistics or flexible plotting of the data. Typically, researchers will only apply more sophisticated statistics and plotting approaches to region of interest (ROI) mean data, but many studies would benefit from tools that enable statistics at voxel-resolution, facilitating the application of almost any R function to voxel-wise TBSS data. Thus, there is dire need for tools that enable users to use any functionality in R, including machine learning applications, to bridge neuroscience and data science.

Here we report the creation and implementation of skiftiTools that bridges this gap for DTI derived scalar maps that have been processed with the TBSS, and the derived skeletons have mapped to standard space template space (supporting Montreal Neurological Institute, MNI templates, or study-specific templates). The skiftiTools functions create standardised and portable tabular data formats and thus allows investigators to benefit from a vast array of statistical tests that are available through R software library, which greatly expands the currently available toolkit for neuroscientists. This alleviates the constraints of the contemporary image-based statistics by allowing many more conventional and advanced statistical approaches, reporting effect sizes and other measures of interest across the whole brain. We provide resources to perform multiple comparison corrections via threshold free cluster enhancement (TFCE) ^11^ that can now be applied to any statistical test that yields p values. skiftiTools is freely available (https://github.com/haanme/skiftiTools), and we provide the tools as a docker container and as a CRAN (The Comprehensive R Archive Network) package repository (https://cran.r-project.org/web/packages/skiftiTools/index.html). In addition, the resources include a thorough documentation with example processing steps (https://skiftitools.readthedocs.io/en/stable/usage.html).

## 2. Overview of skiftiTools and functionality

The skiftiTools writes 3-dimensional voxelwise data into tab-separated values ASCII files, facilitating interoperability with commonly used tools, for example, the R language (RStudio), SPSS, SAS, and GraphPad Prism, among others. One of the main applications is to convert a 4-dimensional subject-per-volume NIfTI into a tabular subject-per-row ASCII. Following statistical processing, the ASCII data that is produced might be read into skiftiTools for visualization of summary statistics and effect sizes. For effective disk usage, the software supports its own stand-alone compressed file format (.skifti), the tab-separated ASCII format, and NIfTI, the most popular MR image image format. Dependencies of the skiftiTools include the following R libraries: *argparse, RNifti, stringr, rmarchingcubes, rgl, abind, png, Rcvg, oce, methods and s2dv*. Transformations between 3D and 4D image volumes and tabular data can enable powerful novel approaches to data modelling as this creates a bridge between neuroimaging software and applications that are available in popular statistical applications of Python, Matlab, and R among others. The transformations ultimately rely on standard mapping of the 3D voxel grid so that the numerical values stored in each voxel are stored to a tabular format in a certain order, creating the possibility to also read the the output from any mathematical manipulation back to the image format and visualised as brain images. The Human Connectome Project was a well-known forerunner in creating such standards for mapping between multimodal volumetric MRI data into tabular format through CIFTI (Connectivity Informatics Technology Initiative) files ^8^. Here, we aimed for a simple as possible application with minimal metadata. The MRI is converted into tabular voxel-per-column data, and due to the standard reading pattern, keeping the data reading and writing standard the metadata are not necessary for easy re-conversion to NifTI. Standardization is necessary as one could read a 3D image file in: up / down + left / right + moving along in x / y / z axis, resulting in 2 x 2 x 3 = 12 different possibilities for vectorizing the data. In addition, voxel coordinates can be saved to facilitate spatially-dependent multiple testing correction of p-values (i.e., TFCE). The NIfTI files are read in slice-by-slice in standard arrangement: x (from Left to Right), y (from Posterior to Anterior), z (from Inferior to Superior) directions, with RNifti package’s readNifti function.

The key functionality of the skiftiTools is summarized in Figure 1. We have created a novel standard for reading in 3D white matter skeletons created with the TBSS, whereby volumetric data is read to a typical tabular format. This tabular data can then be imported to R and used flexibly for analyses. It is important to note here, that we have not included any statistical tests as part of skiftiTools, which was an intentional choice to allow maximum flexibility for users. However, conversion to .skifti format makes the TBSS data compatible with most R functions, enabling a vast range of statistics that would be otherwise impractical or impossible. Use cases are provided below for voxel-wise general linear modelling, non-linear modelling, voxel-wise correlation matrices, principal component analysis, clustering, and elastic net regression predictive modelling. Finally, we describe the computational requirements for all example analyses described below, and have included instructions for running them as part of the online documentation.

**Figure 1.**
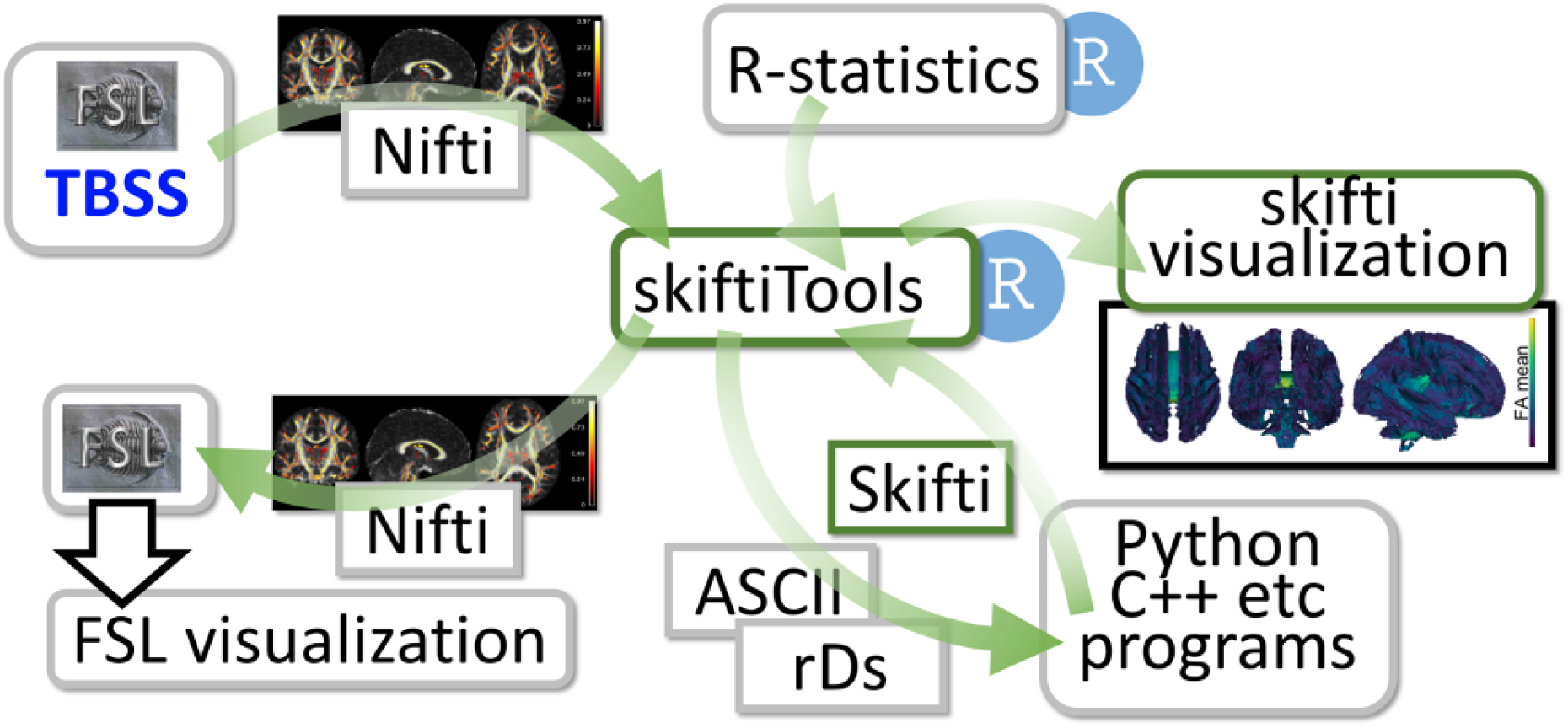
Overview of the skiftiTools functionality.

## 3. Example analyses

To showcase the potential of the skiftiTools, we provide five use case examples: 1) Performing conventional statistical tests (T tests, correlations and linear regression) across the white matter skeleton, and implementation of spatial multiple comparison correction via TFCE that we show to be near identical to those performed with FSL randomize, 2) Mapping non-linear associations between DTI metrics and age in neonates via general additive models, 3) Mapping the voxelwise correlations within the diffusion tensor skeletons to show the joint variability of voxels, 4) Use of dimensionality reductions (principal component analysis [PCA], independent component analysis [ICA]) and clustering (K means clustering, Gaussian Mixture Modeling [GMM]), and 5) Supervised machine learning with elastic net regression to predict postmenstrual age from neonatal scans. Our example data sets are from the beginning of the human lifespan including neonates and 5-year-old children, but the tools can be used in any age group. Effect size maps were converted back to NIfTI using the skiftiTools function Skifti2NIfTI. All other visualisation was done with the commonly used R library *ggplot2*.

### 3.1. Example data sets and DTI image processing

We used preprocessed Developing Human Connectome project data release 3.0 (dHCP study), and the ‘EDDY’ version of the preprocessed DTI data for the purposes of this article ^12^. The other two data sets that we used in the example analyses are from the FinnBrain Birth cohort study (FB study) ^13^; https://sites.utu.fi/finnbrain/en/. The details of the diffusion MRI preprocessing are provided in prior publications for neonates ^14,15^ and for 5-year-olds ^16,17^. All preprocessed data were postprocessed with a pipeline that combined spatial normalization via Advanced Normalization Tools (Tustison 2021) (ANTs; https://stnava.github.io/ANTs/) and TBSS of FSL ^10^ - detailed in (https://github.com/pnlbwh/TBSS). As a result, all images were registered to the ENIGMA (Enhancing Neuro Imaging Genetics Through Meta Analysis) DTI template. The TBSS white matter skeleton contained 117,139 voxels. This produced 4-dimensional NIfTI image files that were used as an input to skiftiTools. All statistical analyses were carried out in R software with variable versions and computers as defined by the computational requirements (see section 3.7. for details on computational requirements). Although we processed the example data in this standardised way for the example data sets, the input can be TBSS data registered to any template space.

### 3.2. Performing conventional statistical tests across the white matter skeleton and implementation of spatial multiple comparison correction

As a first example, we carried out typical statistics in FB data that tested for sex differences, correlations and linear regression of premenstrual age at scan. The novel features that skiftiTools allows include visualisation capabilities that include volcano plots and flexible plotting of effect sizes such as Cohen’s d, correlation coefficients, and regression-derived beta coefficients. Although it is possible in principle to simply extract p-values from the parametric models described above, and adjust these for multiple testing to correct the family-wise error or false discovery rates by traditional methods based on number of tests (e.g., Bonferroni or Benjamini-Hochberg), this is not ideal for MRI data and leads to false negative results. Instead, the corrected p-values should be estimated by permutation testing considering the spatial dependency of the voxels, such as the clustermass or TFCE methods commonly applied to MRI data in NIfTI format (Winkler 2014). We implemented full TFCE correction to skifti data by using the permuco4brain R package (https://github.com/jaromilfrossard/permuco4brain/tree/master) with some important modifications.

Our example analyses show that there were no differences between males and females after TFCE correction, but skiftiTools allowed us to visualise the areas where possible differences could be located by facilitating easy creation of effect size maps (Figure 2A-C). We found multiple brain areas that are positively correlated with premenstrual age at scan (Figure 2D-F). Correspondingly, FA values associated positively with premenstrual age at scan in a regression model that adjusted for sex (Figure 2G-I). Of note, adding sex as a covariate markedly changes the pattern of p values in the volcano plots (Figure 2D vs. Figure 2G). Finally, we confirmed that we were able to replicate results obtained from FSL randomise ^11^, which is the key standard for voxel-wise statistics (Figure 2J-L).

**Figure 2.**
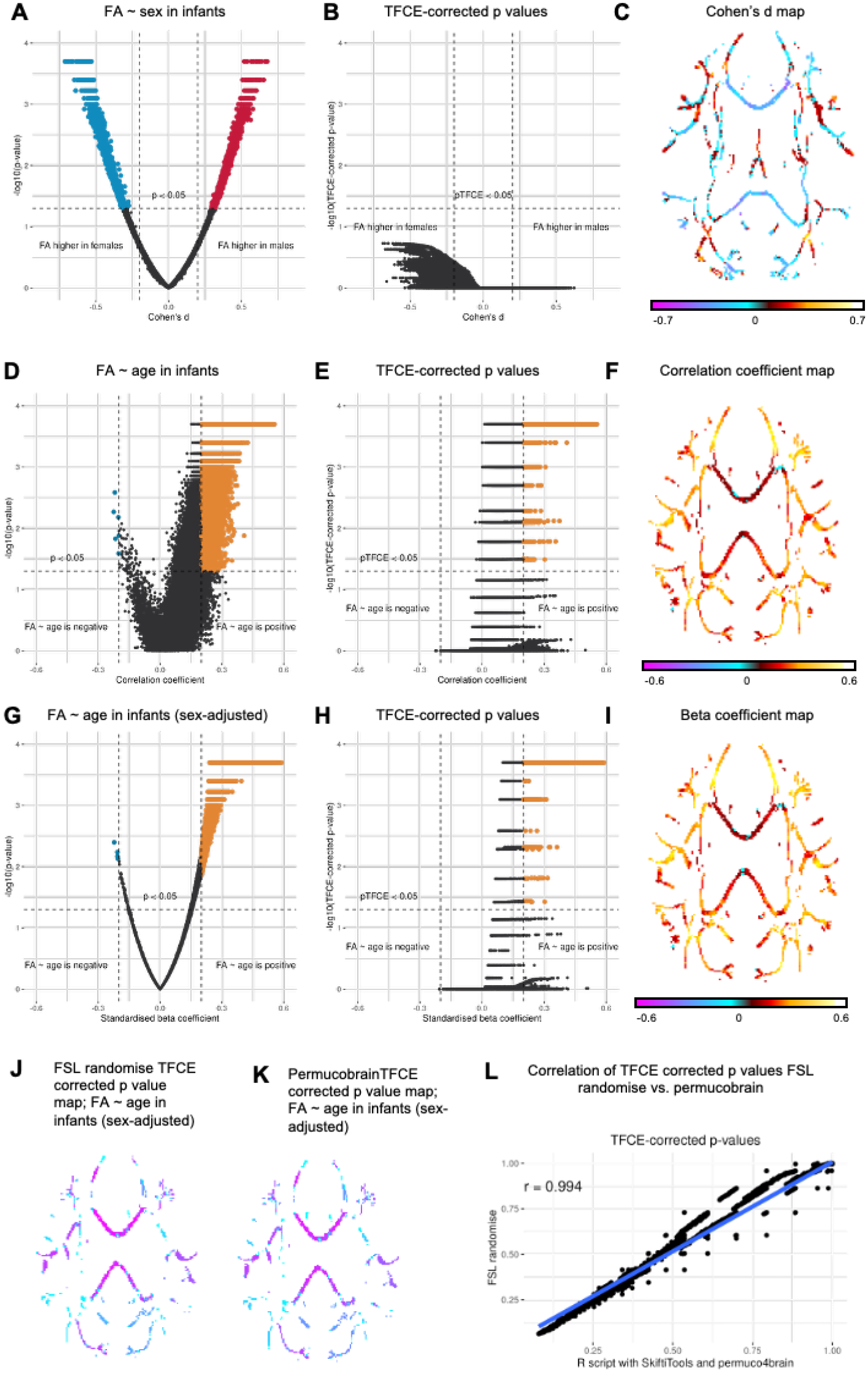
Results showing associations of age and sex to fractional anisotropy (FA) in FinnBrain data sets. A) volcano plot showing uncorrected p values for sex differences, B) related threshold-free cluster enhancement (TFCE) corrected p values, and C) Cohen’s d effect size map visualised on the tract-based spatial statistics (TBSS) skeleton. D) p values from a correlation of premenstrual age and FA, E) TFCE corrected p values, and F) correlation coefficients visualised on the TBSS skeleton. G) Linear regression model showing the association between PMA and FA (adjusted for sex), TFCE corrected p values, and F) standardised regression coefficients visualised on the TBSS skeleton. We replicated the output of FSL randomise (J) with permucobrain-based permutations using tabular data (K), and found that the TFCE corrected p values very tightly associated (L).

### 3.3. Mapping non-linear associations between DTI tensor metrics and age in neonates

It is of high importance to be able to model nonlinear associations of age on brain metrics ^4,18^. Our example analyses fitted standard linear models with R function ‘lm’ and general additive models (GAM) using the ‘mgcv’ library. We fitted linear and nonlinear models to the neonatal dHCP data set. We found that there were significant implications of nonlinear associations in the data (Figure 3), which is highly relevant for all neonatal DTI studies even though further analyses are beyond the scope of the current study.

**Figure 3.**
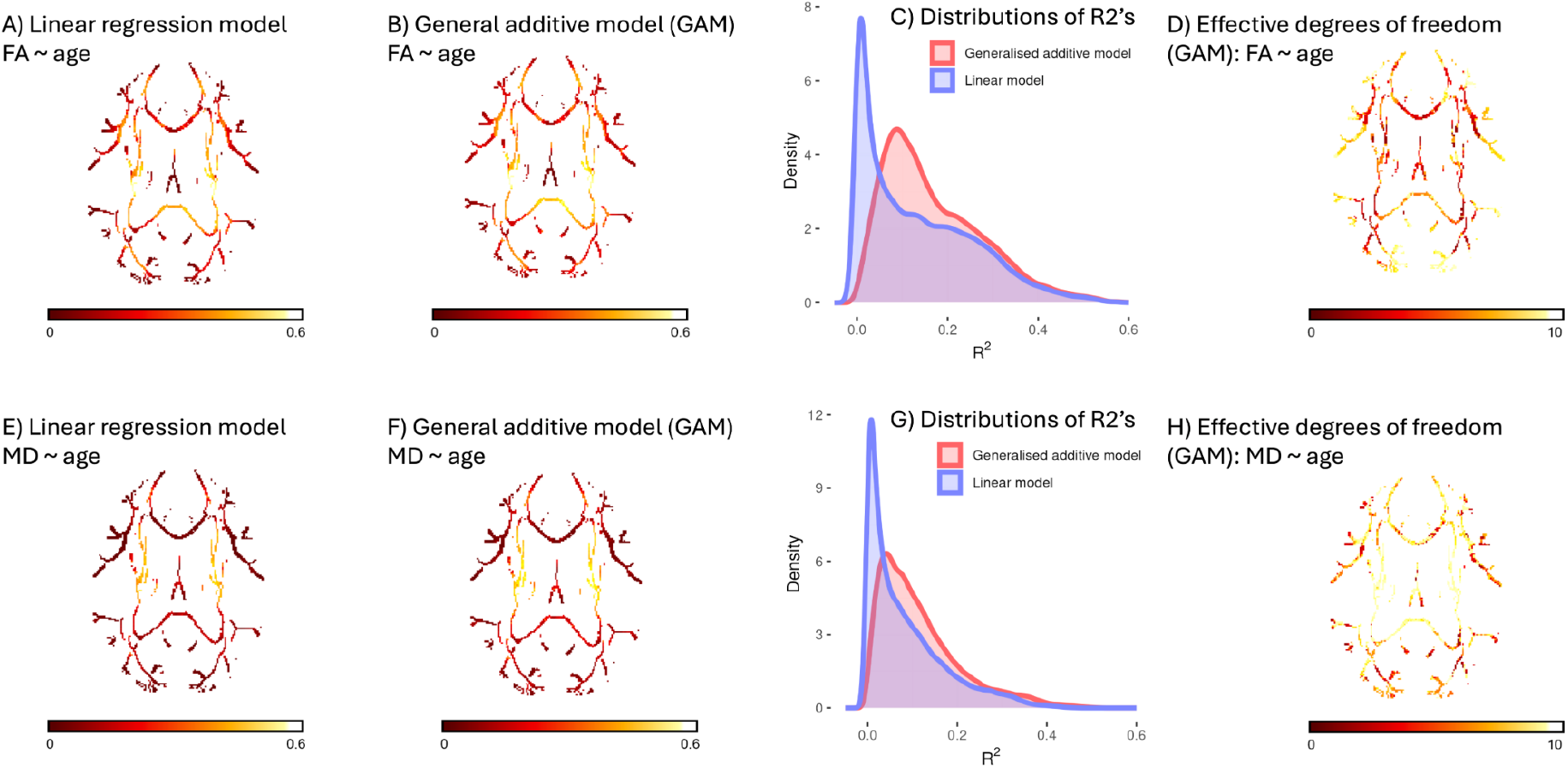
Results showing linear and nonlinear associations of premenstrual age with fractional anisotropy (FA) and mean diffusivity (MD) in the dHCP data set. A) R2 values from the linear model for association between PMA and FA, B) R2 values for a similar nonlinear model, C) differences in voxelwise R2 values between linear and nonlinear models (y axis ∼ n x 1000 voxels), and D) plot of effective degrees of freedom where higher values indicate likely nonlinearity in the associations. E-H) Similar plots for MD values.

### 3.4. Mapping the voxelwise correlations within the diffusion tensor skeletons

We performed voxel-to-voxel correlations of the FA and MD values across the entire TBSS skeleton for all three data sets using R function ‘cor’ (Figure 4). This required high computational capacity (detailed in section 3.7). The correlation plots show a replication of the overall pattern of within-skeleton correlations in neonates (B vs. C), and the developmental changes that are clearly shown as the appearance of negative associations within the skeleton (C vs. D), which is likely indicative of regional differentiation in DTI metrics.

**Figure 4.**
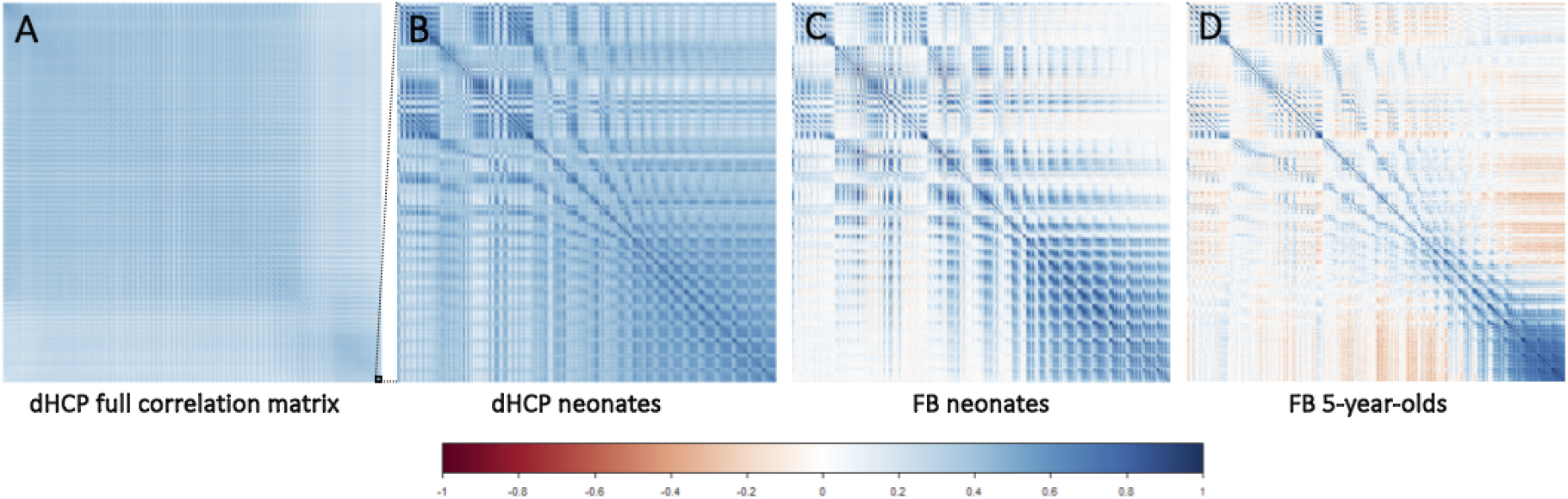
Correlation plots of fractional anisotropy within the white matter skeleton. A) full correlation matrix across 117139 voxels was estimated and the lower portion (marked with a black square in bottom right part of the matrix) was plotted for example figures. B) Partial plot of the correlation matrix estimated from the dHPC neonate data set. C) Partial plot of the correlation matrix estimated from the FB neonate data set. D) Partial plot of the correlation matrix estimated from the FB 5-year-old data set.

### 3.5. Dimension reductions and clustering

Using the neonatal data sets (FB and dHCP), we performed dimension reduction through PCA and ICA as well as clustering with K-means clustering and GMM (Figure 5). The PCA and K means clustering models use base R functions, ICA uses R library ‘ica’, and GMM was conducted with R library ‘mclust’. We found that most PCA and ICA analyses were driven by the first component and that the results overall were variable between the data sets. In the FB data set (plotted here as an example), we found that the first principal component did not vary between males and females and that it had a strong negative association to gestational age (ß = -0.70, p < 0.0001) (Figure 5). K means clustering provided separate sub groups with three clusters (K = 3) for FA, but the clustering results for MD were less distinct (Figure 5).

**Figure 5.**
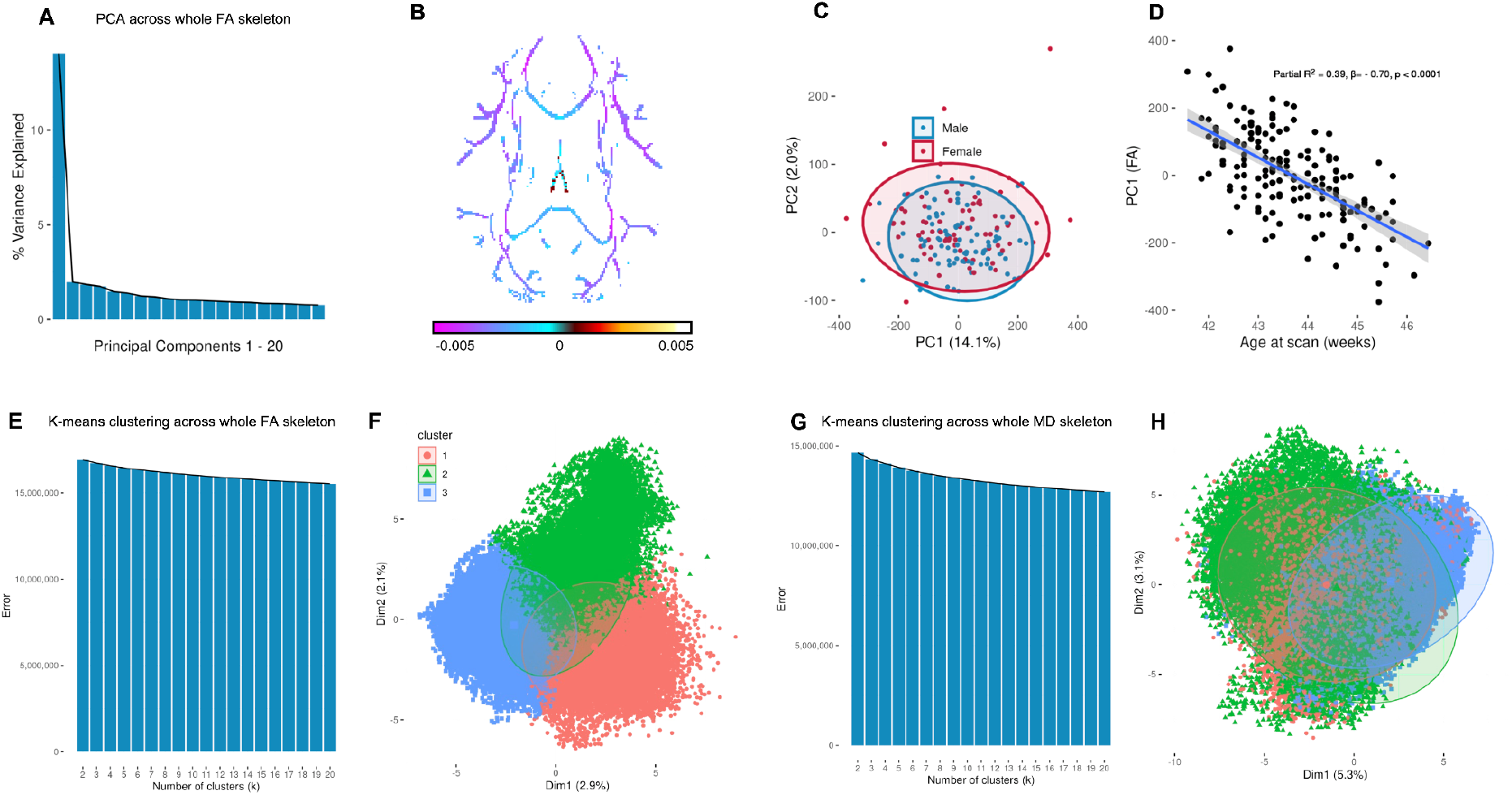
Dimension reductions and clustering analyses of the white matter skeleton across 117139 voxels. A) Principal component analysis (PCA) of fractional anisotropy, and B) related visualisation of first principal component weights in the brain. C) PCA components did not differ between males and females, but D) showed a strong association with pre-menstrual age at scan. E-F) K means clustering across the fractional anisotropy values yielded distinct clusters, but G-H) mean diffusivity values did not form clear clusters in K means clustering.

### 3.6. Elastic net regression to predict postmenstrual age in neonates

We applied five-fold cross-validated elastic net regression to predict postmenstrual age of the neonate samples (FB and dHCP), with the entire TBSS skeleton as the prediction matrix. We used a recently published R library ‘biglasso’ that enabled very efficient computations to run the analyses. In the dHCP data, the elastic net regression achieved a very good prediction (average R2 = 0.83 [+/-0.03], normalized root mean squared error; NRSME = 0.08 [+/-0.005]). In the FB data, the prediction did not perform equally well (average R2 = 0.36 [+/-0.03], NRSME = 0.21 [+/-0.01]), most likely due to smaller sample size, smaller age range, and lower variability between PMA and MRI scan. Interestingly, the voxels contributing to the predictions were scattered across the brain and we observed both positive and negative coefficients (Figure 6).

**Figure 6.**
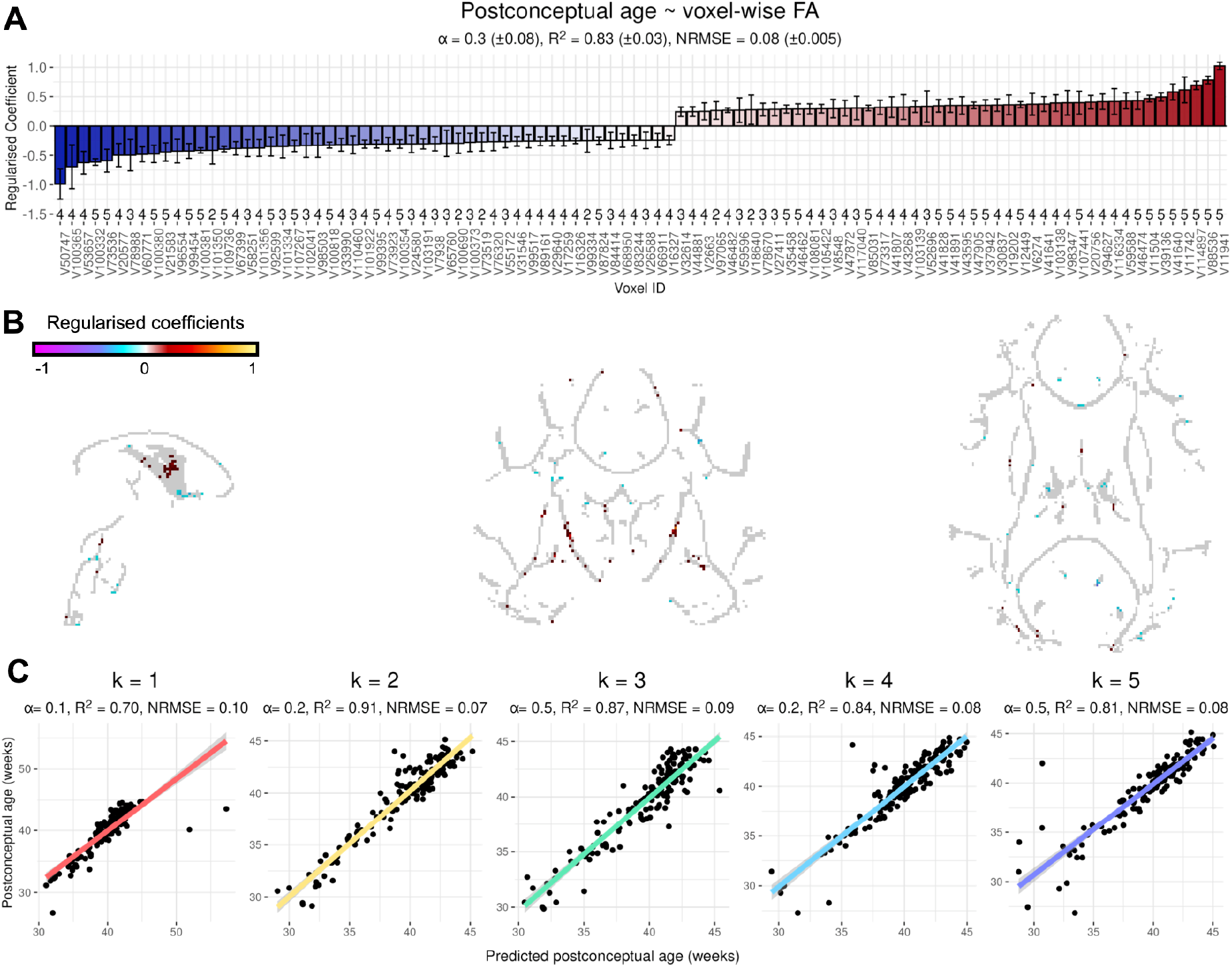
Elastic net regression results from the dHCP data set. Predictions of premenstrual age at scan across all of the white matter skeleton across ∼ 192k voxels. A) Voxels with largest coefficient are plotted, and visualised on the white matter skeleton in B. C) predictions from k = 1–5 folds.

### 3.7. Computational requirements

After reading the skifti file into an R session, this is simply a dataframe where each column represents the signal intensity (e.g., FA) in a given voxel. Thus, using the ENIGMA DTI skeleton template, as in the example analyses, there will be 117,139 columns plus one additional column for subject IDs.

- **Sections 3.2 & 3.3. conventional statistics:** For T tests, linear models, and non-linear models one can simply loop a statistical model over each column/voxel and extract the parameters of interest into a dataframe. This is not computationally demanding and will run in a few hours on a typical computer without the need for parallelisation.
- **Section 3.2. Spatial multiple comparisons correction with TFCE:** Unlike looping a simple model over voxels, the brainperm function is extremely computationally expensive with 5000 permutations. For example, when applied to a dataset of 117,139 voxels and 112 participants, the model requires 30–40 GB of random access memory (RAM). The function supports easy parallelisation with the ‘futures’ R package, and loads it as a dependency. This is convenient as long as your system has enough total RAM memory to run the model. For instance, a model distributed over 6 worker processes with 5000 permutations of a two-tailed test will take 2 to 3 hours. We ran the analyses in CSC’s Puhti supercomputer cluster (https://csc.fi/).
- **Section 3.4. estimation of voxel-to-voxel correlation matrices:** The ASCII skifti files ranged in size from 310 MB (FinnBrain 5-year-old sample, N = 147) to 1.7 GB (dHCP neonatal sample, N = 778). After reading the file into an R session, the correlation matrix is estimated directly. It is not practical to estimate the correlation matrix across the entire dataset using cor(skifti), because this involves 2n calculations, where n is the number of columns/voxels. Thus, performing 2,117,139 calculations at a rate of one per second would take approximately 2,117,139 seconds (2,117,139 ÷ 60 = 35,285.65 minutes; 35,285.65 ÷ 60 = 588.09 hours; 588.09 ÷ 24 = ∼24.5 days) to complete. To make the computations faster, it should be done on a per-column basis in a for-loop, and there will instead be only n operations, so at 1 second per calculation, this would take 32 hours. We ran the analyses in CSC’s Puhti supercomputer cluster (https://csc.fi/) due to considerable computational requirements that could only partly be solved by parallelization. For instance, holding the correlation matrix in memory requires 100 GB RAM, while plotting it uses approximately another 500 GB RAM and takes 2–3 hours.
- **Section 3.5. Dimension reduction and clustering:** PCA and ICA did not pose computational challenges. The computing resources required for K-means and clus-tering vary considerably depending on your sample size, but it is unlikely that one would be able to run it on a standard computer. Thus, we ran the analyses in CSC’s Puhti supercomputer cluster (https://csc.fi/), using foreach and the parallel backend doMPI. Parallelizing over 19 values of k achieved a roughly 12.5-fold speed-up (total time required was 3 hours for dHCP neonatal and 45 minutes for FinnBrain infant datasets, using up to 10 GB memory per CPU). For Gaussian Mixture Modeling, parallelization achieved almost perfect speed-up (18-fold over 20 parallel tasks, taking 2.5 hours with 10 - 15 GB memory per CPU for dHCP neonatal, and 30 minutes for FinnBrain neonatal with 2 - 5 GB memory per CPU).
- **Section 3.6. Elastic net regression:** Tabular MRI data in the form of a skifti file allows for full voxel-wise supervised machine learning models, such as elastic net (including ridge and LASSO) regression. We used the R package biglasso, which stores data on disk and only reads it into memory when necessary during model fitting. To fit an elastic net regression using all 117,139 voxels from a TBSS skeleton as predictors, on datasets with 169 or 778 participants, the total memory requirements were only 2 GB and 5 GB, respectively. A single 80:20 train/test split with a predefined α level (mixture parameter to define the type of penalty applied) and nested 10-fold cross validation to choose λ (the magnitude of the penalty applied) over a grid of 1000 possible values takes 15 minutes or 2 hours for the small and large datasets, respectively. Thus, the model can be run on any system without need for parallelisation, but naturally if the sample size is scaled up it may take much longer to run.

## 4. Discussion

Here, we describe a novel tool that allows conversions between NIfTIi images and tabular data format for the white matter tract centre estimates (white matter skeleton) that is derived from FSL’s TBSS. This work provides a powerful and flexible tool for storing and analyzing complex brain imaging data in standard tabular format, facilitating advanced neuroimaging research and analysis. The example analyses were carried out in the MNI template space and the white matter skeleton of the ENIGMA consortium, but the tools allow users to specify data-specific template spaces, and the functionalities can be extended used for any binary 3D volume in any template space in the future. We have named the tabular data format of the TBSS skeleton as *skifti* files based on Scandinavian word *skifta* (translating to change, exchange, divide, distribute). The Faroese base version of the word *‘skifti’* was chosen as the final name as the ending matches those used in other high dimensional data types such as *gifti* and *CIFTI* files.

A growing number of neuroimaging tools are being developed to enable data analysis in a more tool-independent manner when analysing three- and four-dimensional brain imaging data in various statistical settings. Ideally, such tools still offer user interfaces that are accessible to those with a basic understanding of programming and statistics. The traditional methods for doing statistical computations in voxel-space for brain DTI data in brain image processing are TBSS and Statistical Parametric Mapping (SPM). Although these tools contain methods for fundamental, general purpose statistical analysis through general linear models (GLM), the focus on GLM-based statistics, p value – based results reporting that omits effect sizes, and lack of support for commonly used non-linear models are a major limitation. They might also not be readily scalable for application to larger datasets and integration of advanced statistical methods such as machine learning, nor are they able to support meta analyses across voxelwise summary measures.

We identified only one existing R software application tailored for diffusion MRI data. The ModelArray is a python toolbox that is operated together with R ^4^. Despite the possibility to create custom functionality, the ModelArray is optimized to support only two main statistical tests, i.e., linear models as well as generalized additive models and has impressive optimization for the use of computational resources. It is important to note that ModelArray can be tailored to read in fixelwise, voxelwise, and CIFTI format data to perform flexible analyses through linear or non-linear models and enables the investigators to visualise the results. The skiftiTools is complementary to ModelArray, but offers significantly more flexibility for statistical tests. We note that all statistical tests were carried out with existing R libraries that are not part of the skiftiTools. Finally, prior work has not introduced possibilities to carry out spatial multiple comparison corrections ^4^, which we argue is a key feature to bridge the novel tools of DTI statistics to mainstream neuroimaging software such as FSL ^19^.

The major limitation of skiftiTools is related to the large tabular data size that necessitates the R libraries to support such data dimensions. For instance, the creation of correlation matrices required considerable computational resources, and such analyses were only possible in high-performance computing environments. Optimizing R libraries is beyond the scope of this article, and we would like to note that most analyses, including voxel-resolution elastic net regression, were easily implemented with typical computational resources, i.e., when the native R libraries support large data dimensions, the analyses run smoothly.

In conclusion, the lack of support for basic and advanced statistical operations forces users to use and report predominantly probabilistic voxel-wise statistics or limit their research to region of interest analyses. skiftiTools enables investigators to carry out analyses that are not available in mainstream software such as the calculation of effect sizes, plotting options, provides full support for spatial multiple comparisons via permutation testing, and enables use of machine learning libraries. The results of the analyses can be directly visualised with skiftiTools and saved to NIfTI files for visualisation in any software of choice.

## Data Availability Statement

The developing Human Connectome Project data can be found in online repositories and are openly available. The data used in this study stemmed from the third data release: http://www.developingconnectome.org/data-release/third-data-release, and the most recent data is shared in NDA data base: https://nda.nih.gov/edit_collection.html?id=3955. The data stemming from FinnBrain Birth cohort study data cannot be shared openly due to The Finnish law and ethical permissions. Data sharing is possible via formal agreements and interested investigators are requested to contact FinnBrain study administration; (https://sites.utu.fi/finnbrain/en/contact/).

## Code availability

skiftiTools is freely available (https://github.com/haanme/skiftiTools), and we provide the tools as a docker container and as a CRAN (The Comprehensive R Archive Network) package repository (https://cran.r-project.org/web/packages/skiftiTools/index.html). In addition, the resources include a thorough documentation with example processing steps (https://skiftitools.readthedocs.io/en/stable/usage.html).

## Acknowledgements

The authors wish to acknowledge CSC – IT Center for Science, Finland, for computational resources.

## Author contributions

- JJT: conceptualization of the study, tools and functionality, editing and testing code and implementing the postprocessing pipelines, performing and supervision of formal statistical analyses (by AB), leading the manuscript writing.
- AB: Carrying out the example analyses and implementing the necessary high-performance computations, and creation of TFCE correction pipeline.
- AJ: configured and tested the Docker environment, contributed to pipeline documentation
- IS: Submitting the R package to CRAN
- HKA: compiled and deployed the pipeline documentation
- AR: contributed to pipeline documentation
- IMW: checked the R code base, contributed to pipeline documentation
- EV: manuscript writing
- SL: manuscript writing
- EPP: manuscript writing,tested Docker environment
- HK: manuscript writing
- RK: manuscript writing
- LK: manuscript writing
- AA: manuscript writing
- JS: manuscript writing
- RAIB: manuscript writing
- HM: Conceptualization of the study, tools and functionality, writing R code and implementing the skiftiTools, supervision of software development project, manuscript writing. **All authors critically reviewed the manuscript and accepted it in its final form prior to submission**.

